# A novel coronavirus associated with enteritis in broiler chickens, France, 2025

**DOI:** 10.64898/2026.06.09.731082

**Authors:** Mattias Delpont, Nicolas Gaide, Vincent Blondel, Manuela Crispo, Benjamin Linard, Aurélie Sécula, Mathilda Walch, Lilou Bortot, Elodie Durand, Isabelle Fourquaux, Léni Corrand, Sébastien Soubies, Pierre Bessière, Guillaume Croville, Jean-Luc Guérin

## Abstract

Coronaviruses of the genus *Gammacoronavirus* cause major poultry diseases, including infectious bronchitis in chickens and enteritis in turkeys and guinea fowl. Until now, no enteric coronavirus distinct from infectious bronchitis virus had been reported in chickens. Between late 2024 and 2025, severe enteritis outbreaks affected broiler farms in southwestern France, causing increased mortality, wet litter, cyanosis, lethargy, ruffled feathers, and high slaughter condemnation rates. Necropsy and histopathology revealed diffuse enteritis and dehydration. Metagenomic sequencing identified abundant coronavirus reads as the only pathogenic viral signal. Whole-genome phylogeny showed a novel gammacoronavirus lineage closely related to guinea fowl coronavirus but distinct from infectious bronchitis virus and turkey coronavirus. Viral RNA was detected in enterocytes by RNAscope *in situ* hybridization, and electron microscopy revealed coronavirus-like particles. These findings describe a novel enteritis-associated coronavirus in broiler chickens (ChECoV), although the drivers of its host-range expansion into chickens remain to be elucidated.

## Introduction

Poultry production is a cornerstone of global food security, with chicken meat representing one of the most consumed sources of animal protein worldwide (1). The sustainability of this sector depends heavily on the control of infectious diseases that can rapidly spread through intensive and free-range systems alike. Among these, coronaviruses are of particular concern given their capacity for cross-species transmission and rapid genetic diversification.

Coronaviruses of the genus *Gammacoronavirus* are recognized causes of disease in poultry. While infectious bronchitis virus (IBV) is typically associated with respiratory, renal and reproductive conditions, infection with turkey coronavirus (TCoV) and guinea fowl coronavirus (GfCoV) can result in enteritis, leading to severe growth retardation and massive mortality (2–5). So far, outbreaks of clinical enteritis involving GfCoV have been sporadic, and only reported in guinea fowl (*Numida meleagris*) in France. To date, no reservoir species were identified (6–8).

Enteric diseases in poultry are usually considered challenging and complex conditions, due to the involvement of multiple etiological agents (co-infections, including viruses, bacteria and parasites). Their clinical presentation is highly dependent on the production system and environmental factors and they are usually described at a rather young age (9,10).

Between late 2024 and 2025, a series of outbreaks of enteric disease was recorded in free-range broiler farms in south-western France. The affected chickens were approaching the end of their production cycle. The disease was characterized by unusually high mortality rates and severe intestinal lesions. Considering the atypical presentation and the severity of these outbreaks, an in-depth diagnostic investigation was conducted to characterize the disease and identify potential causative agents involved.

## Materials and Methods

### Selection of cases and sampling

After initial reports in late 2024, four outbreaks investigated between February and June 2025 were included in this study. The 4 farms were 2 to 120 km apart, with no identified epidemiological link between them. Farms were visited within one week after the appearance of clinical signs. Farm records, when available, were collected. For each flock, 20 cloacal swabs were randomly taken from live birds at the farm and selected animals, including freshly dead and clinically-affected, euthanized birds, were sent to the laboratory for diagnostic investigation. Environmental wipes were also taken to collect dust from barn walls, feeder and drinker lines in some of the farms. A complete necropsy was performed on all subjects and lesions were recorded. Swabs, intestinal contents and tissue sections were sampled and processed for molecular and histopathological analysis.

### Histopathology

Tissue sections collected at necropsy from representative chickens belonging to the four cases: 59/2025 (n=4 birds), 93/2025 (n=4), 239/2025 (n=8) and 295/2025 (n=6), were fixed in 10% neutral buffered formalin, trimmed, paraffin-embedded and cut at 3 μm thickness. Selected tissues included small intestine, pancreas, kidney, spleen (n=22 birds for each organ), bursa (n=21), proventriculus (n=16), thymus (n=14), trachea, lung and gizzard (n=12). Sections were routinely stained with hematoxylin and eosin (H&E) for histopathological assessment by light microscopy, and scanned at x20 magnification using an Olympus Slideview VS200 scanner.

### Nucleic acid extraction and PCR

Swabs, tissues and environmental samples were submitted to molecular analysis. Nucleic acids were extracted and amplified as follows: viral RNA was extracted from intestinal contents using the ID Gene Mag Fast Extraction Kit on an IDEAL 32 automate (Innovative Diagnostics, Grabels, France), and wipes were processed according to Filaire and colleagues (11). For each clinical case, a real-time RT-PCR targeting gammacoronaviruses was performed on intestinal samples (intestinal contents or cloacal swabs) and wipes (12).

### Metagenomics workflow

For metagenomic analysis, random amplification of RNA was carried out following the SMART9N protocol (13), after which sequencing libraries were prepared using the Native Barcoding Kit 24 V14 (Oxford Nanopore Technologies, Oxford, UK) with nucleic acids from the four clinical cases multiplexed into a single library. The library was loaded onto a FLO-PRO114M flowcell on a PromethION 2 Solo using MinKNOW v24.11.10 and sequenced for 61 hours, with raw data basecalled in real time using Dorado v7.6.8 and a minimum Q score of 10.

The FASTQ files were processed as follows: first, a size filtering with a minimum length required of 500 bp and 30 bp trimming on both ends of the reads using fastplong v 0.2.2 (14,15). The filtered data was then aligned to the *Gallus gallus* genome (NCBI RefSeq assembly GCF_016699485.2) with minimap2 v 2.28 (16,17) and the Samtools package v 1.20 (18) to remove host sequences from the data. Finally, a taxonomic sequence classification was done with Kraken2 v 2.1.2 (19) using the RefSeq complete archaea_bacteria_viral database. The metagenomics classification was visualized using Pavian (20).

The most relevant viral species were selected among the taxonomic classification and used as reference genomes for alignment of sequences with minimap2 and the Samtools suite. Finally, the alignment with the best mapping results (in terms of genome coverage) was chosen to generate a consensus sequence with iVar (21). The alignment was visualized with IGV v 2.17.4 (22) to assess its quality and to identify potentially variable regions.

### Phylogeny and recombination analysis

Sequence alignment was performed using MAFFT online (23), followed by phylogenetic analysis using the maximum likelihood method with IQTREE (24) with the following parameters: bootstrap branch support: ultrafast 1,000 replicates, GTR+F+R4 model (complete genomes) and bootstrap branch support ultrafast 1,000 replicates, TIM3+F+G4 model (spike). Simplot v. 3.5.1 was used to identify potential recombination events (25). Details on SHERPAS recombination search methods are provided in Supplementary material (S1).

### RNAscope *in situ* hybridization

RNAscope *in situ* hybridization (ISH) was performed on a total of nine birds from three cases: 59/2025 (n=3), 239/2025 (n=3), and 295/2025 (n=3). Examined tissues included small intestine, pancreas, thymus and spleen (n=9); kidney and bursa (n=8); caecum (n=6); and trachea, proventriculus and lung (n=3). The probe used in the assay was designed to target the avian coronavirus nucleoprotein sequence from 715-1191 of KF996286.1 which aligned with the detected ChECoV from metagenomic data. Tissue sections (3μm thick) were obtained from formalin-fixed, paraffin-embedded tissue blocks and mounted on positively charged glass slides.

The RNAscope assay was performed using the RNAscope 2.5 HD RED kit according to manufacturer’s instructions. Mild, moderate and extended permeabilization conditions were evaluated and the extended permeabilization protocol was selected based on optimal signal quality. DapB probe was used as a negative control. A confirmed case of influenza encephalitis and a M Gene avian influenza A virus probe served as control to validate the assay technique (26).

### Transmission electron microscopy

Ultrastructural analysis was performed on three chickens belonging to the same flock (239/2025). Animals were selected based on the degree of autolytic changes, the presence of representative histopathological lesions, and detection of viral RNA within tissue sections. Transmission electron microscopy methods are detailed in Supplementary material (S1).

## Results

### Clinical and pathological findings

Affected broilers were slow-growing, free-range chickens, typically processed after 81 days of age. Increased mortality, up to 7% over the course of a week or two, and clinical signs were observed starting usually around 60 [51-70] days of age (Suppl. material – S2). Clinically-affected birds showed mainly cyanosis of the comb and wattles (Figure 1A) associated with lethargy, ruffled feathers at the base of the neck and occasionally tremors and pasted vents. Morbidity, as estimated during farm visits, varied between 0.5% and 5%. Overall condemnations rates at the processing plant ranged from 0.6% (normal) to 2.5%, with most condemnations attributed to a dark red skin coloration of the carcasses. One flock reached 7.5% of downgrades (Suppl. material– S3).

**Figure 1.**
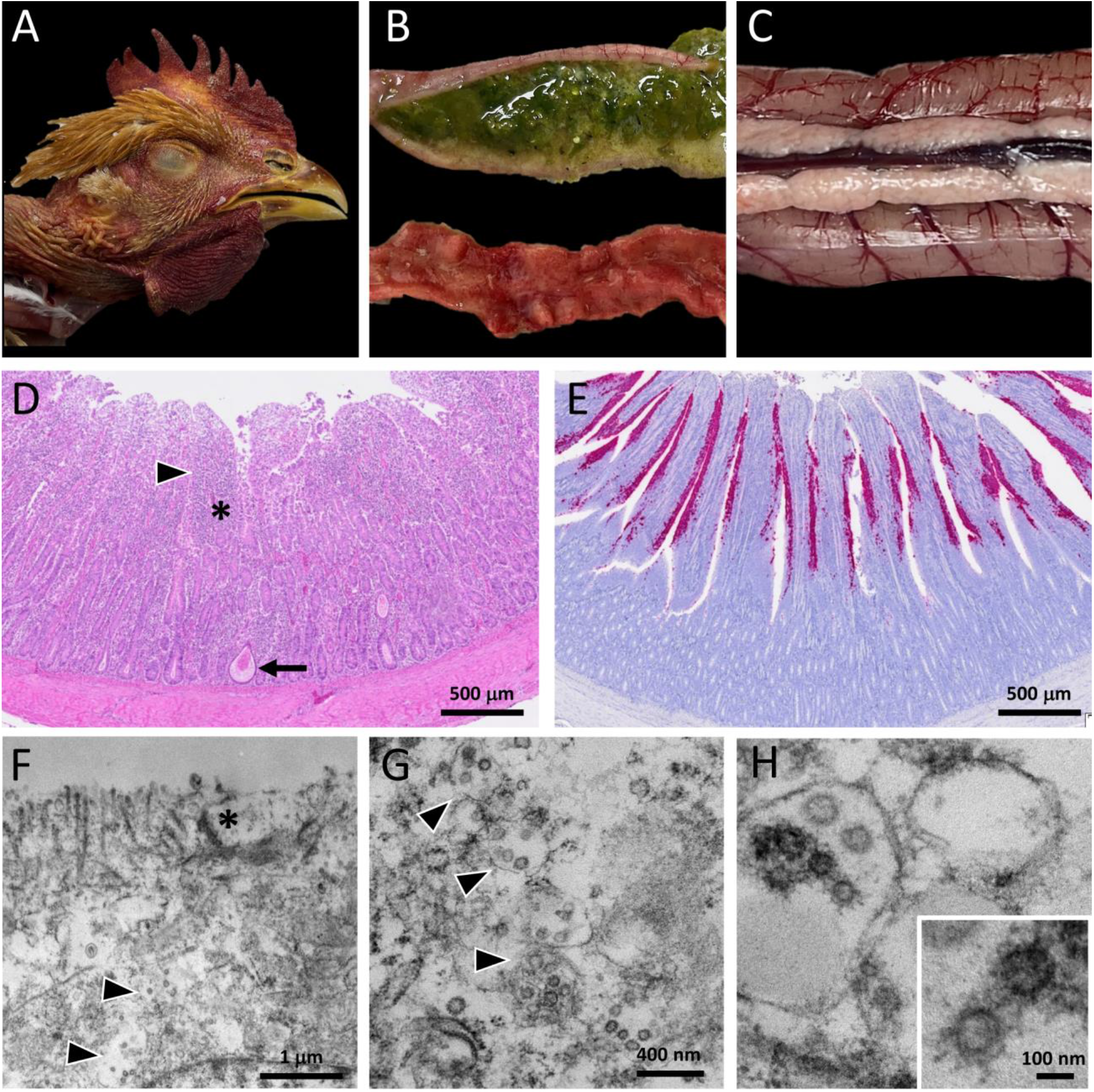
Pathological and ultrastructural findings in chickens naturally-infected with ChECoV. **A**, Head, dark red discoloration of wattle and comb. **B**, Small intestine, the mucosa is thickened with mucoid and watery content (up), or diffusely red (down). **C**, Pancreas, pancreatic lobes are mildly granular with a “snowy” appearance. **D**, Duodenum, the mucosa exhibits villous atrophy, fusion and blunting (arrowhead). The lamina propria is infiltrated by leukocytes (asterisk). Mucosal glands exhibit multifocal segmental dilatation with epithelial attenuation and epithelial sloughing (arrow). Hematoxylin and eosin stain (H&E) and scale bar (sb) = 500 μm. **E**, Duodenum, viral RNA detection is positive and widespread along villous epithelium. RNAscope in situ hybridization targeting coronaviral nucleoprotein RNAs. sb= 500 μm. F, Small intestine, enterocytes in the villous epithelium have disrupted, irregular apical brush border (asterisk). Enterocyte cytoplasm is pale, homogeneous, with distended vesicles containing round particles (arrowheads). Transmission Electron Microscopy (TEM). Bar, 1μm. G and H, Small intestine, numerous distended vesicles (arrowheads) containing several round spherical 90-100nm particles with a biomembrane, heterogenous electron-dense content (insert), peripheral surface projections. TEM. Bar, 400 nm (G) and 100 nm (H).

At necropsy, intestinal contents appeared watery, pale or red and admixed with undigested feed particles, mucoid or absent (Figure 1B). Intestinal walls were frequently thickened due to the presence of pseudomembranes covering the mucosal surface. The pancreas commonly exhibited a granular, “snowy” appearance (Figure 1C). Other remarkable findings included dark red-brown comb and wattles, diffusely dark and dry skeletal muscles, pale and swollen kidneys with urate deposits within the ureters. In two cases multifocal hemorrhages were noticed in the proventricular mucosa. Crop and gizzard impaction with fibrous material (feathers, and bedding - straw) was also frequently observed.

Histopathology revealed diffuse villous atrophy, fusion and blunting, associated with necrosis of villous enterocytes and lymphocytic infiltration of the lamina propria (86% of birds), as well as multifocal dilatation of intestinal crypts (Figure 1D). The second most consistent lesion was extensive pancreatic acinar atrophy with depletion of zymogen granules (86%). Less frequent findings included diffuse thymic and bursal atrophy, multifocal gizzard erosions, focal to extensive proventricular necrosis, lymphocytic tracheitis, exudative bronchitis, and rare foci of renal tubular mineralization (Suppl. material – S4).

### Molecular detection and sequencing

Coronaviral sequences were detected in all four chicken samples, accounting for 5, 80, 81 and 90% of viral reads. No reads corresponding to avian influenza or Newcastle disease viruses, or any other significant avian virus, were detected. No bacterial sequences provided a suitable etiologic candidate for enteritis-associated disease. Complete metagenomics data for viral and bacterial species are provided in Suppl. material (S5). The Sankey diagram presenting the viral and the bacterial content of the metagenomics analysis run on sample ChECoV/S59/2025 is provided in Suppl. material (S6). All included cases presented positive results for gammacoronavirus, on different types of matrices. Most samples presented a very high viral load, with Ct values frequently below 20. PCR data are detailed in Suppl. material (S7).

The strains were later referred to according the recommendations for a Standardized AvCov nomenclature (27) and the corresponding whole-genome sequences are available in GenBank under the accession numbers PV843471.1 (GammaCov/AvCov/chicken/France/S59/2025), PV843472.1 (GammaCov/AvCov/chicken/France/S93/2025), PX987256.1 (GammaCov/AvCov/chicken/France/S239/2025) and PV843474.1 (GammaCov/AvCov/chicken/France/S295/2025). They will be referred hereafter to under a simplified, vernacular terminology Chicken enteritis coronavirus (ChECoV), in consistence with the other poultry AvCoV.

### Genome organization, phylogeny and recombination analysis

The four genomes ranged in size from 27,553 to 27,762 nucleotides, excluding the poly(A) tails. Four of the five genomes contained 13 genes (pp1a–pp1b–S–3a–3b–E–M–4b–4c–5a–5b–N–6b), whereas the 6b gene was absent in one genome (ChECoV/S239/2025). This gene is part of a cassette of accessory genes.

Based on phylogeny analysis conducted on the whole genome and spike protein gene, both ChECoV, GfCoV (recent and older French strains) clustered together (Figure 2). The Simplot analysis did not suggest major recombination events but rather a highly significant genetic divergence among avian coronaviruses at the level of the spike gene; In particular, GfCoVs showed a very significant evolution between 2011, 2017, and 2025 (Figure 3). In the same way, the recombination analysis with SHERPAS did not show any recombination signal in the spike region, that was entirely assigned to the GfCoV type. Notably, a putative recombination was detected in the Orf-1a, PL-Pro region (from positions 5 to 8kb) from the ChECoV/S59/2025, ChECoV/S93/2025 and ChECoV/S295/2025 genomes with assignation to the IBV-QX like type, which may indicate a more ancestral event (Suppl. material – S8).

**Figure 2.**
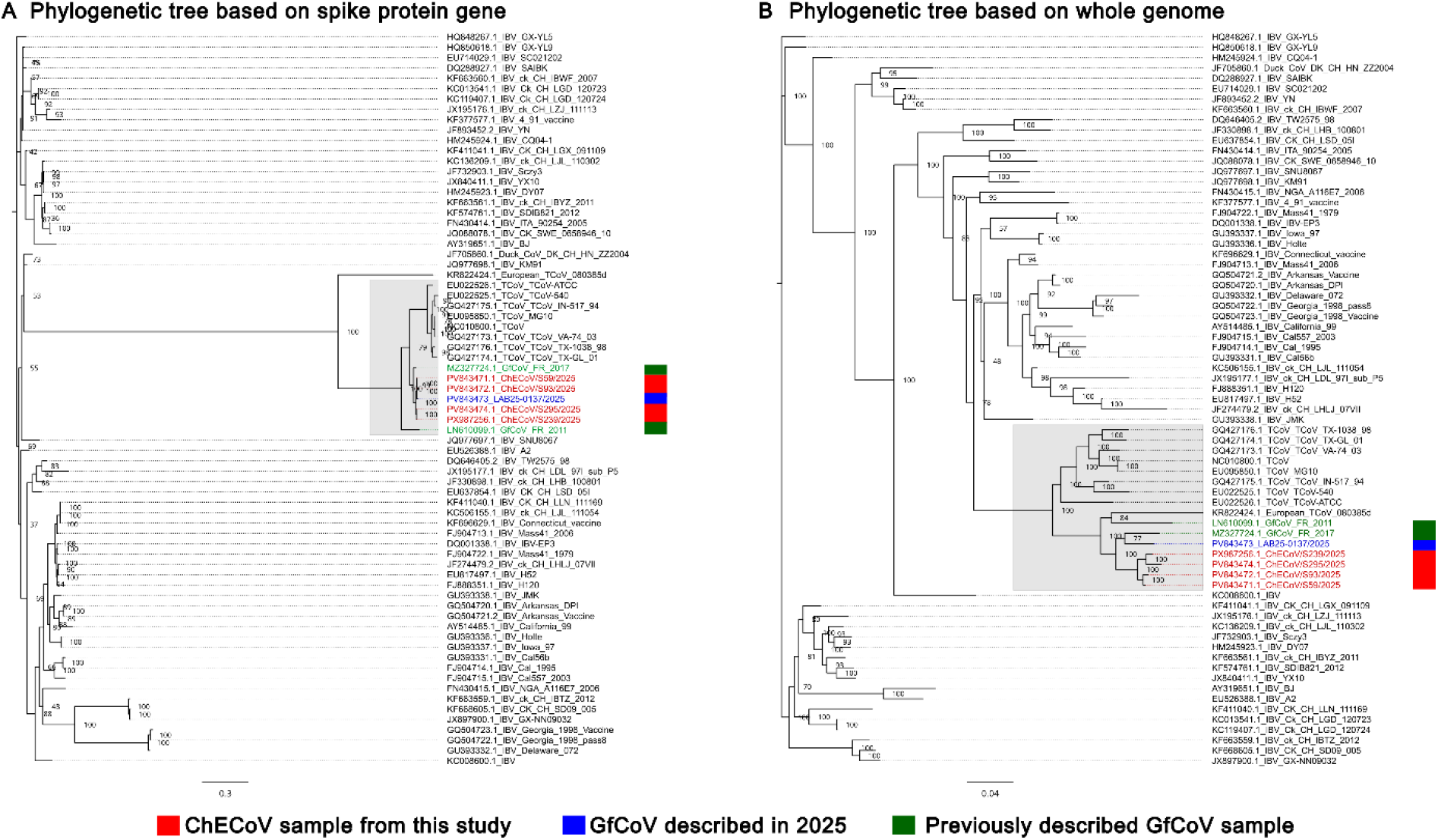
Phylogenetic analysis of Avian CoV spike nucleotide sequence (A) and whole Avian CoV genomes (B). The nucleotide sequences of 7 isolates of chicken or guinea fowl CoV were aligned using MAFFT with representatives of IBV and TCoV lineages. A maximum-likelihood tree was generated based on the alignment. The guinea fowl CoV strains from 2011 and 2017 are colored in green while the chicken and guinea fowl CoV strains of this study are colored in red and blue, respectively. Numbers at nodes indicate bootstrap support (1,000 replicates), and the scale bar indicates the estimated number of substitutions per site.

**Figure 3.**
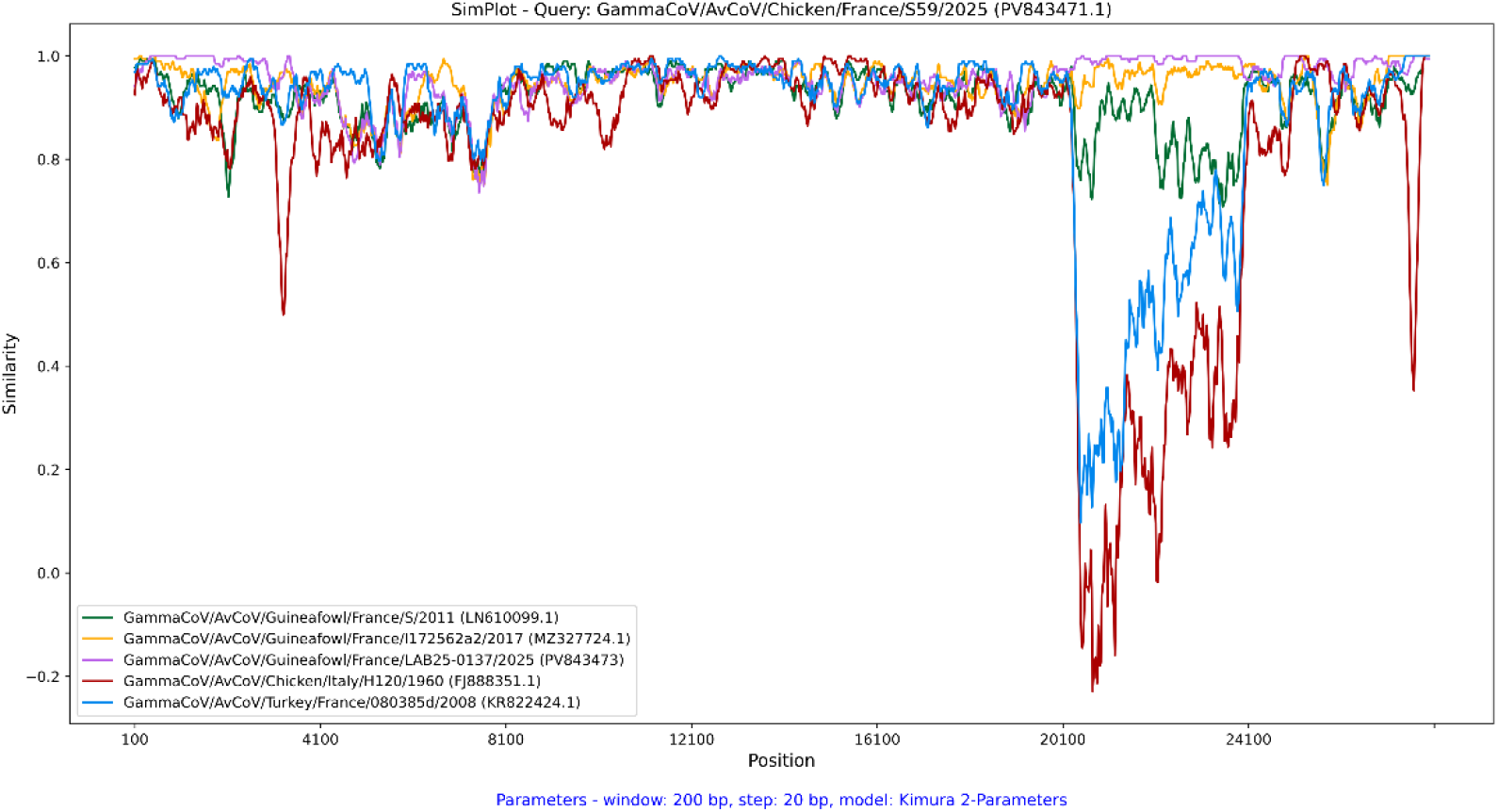
Sliding-window similarity analysis of avian gammacoronavirus genomes. Nucleotide similarity profiles generated using SimPlot for GammaCoV/AvCoV/Chicken/France/S59/2025 (PV843471.1) as the query sequence, compared with representative avian gammacoronaviruses: GammaCoV/AvCoV/Guineafowl/France/S/2011 (LN610099.1), GammaCoV/AvCoV/Guineafowl/France/I172562a2/2017 (MZ327724.1), GammaCoV/AvCoV/Guineafowl/France/LAB25-0137/2025 (PV843473), GammaCoV/AvCoV/Chicken/Italy/H120/1960 (FJ888351.1), and GammaCoV/AvCoV/Turkey/France/080385d/2008 (KR822424.1). Similarity values were calculated using the Kimura two-parameter (K2P) model with a 200-bp sliding window and a 20-bp step size. The x-axis represents nucleotide positions along the viral genome, while the y-axis indicates the level of sequence similarity between the query and reference genomes. Regions displaying pronounced decreases in similarity, particularly between positions ∼20,000 and 24,000 corresponding to the spike (S) gene, indicate substantial genetic divergence among the analyzed strains. Colors correspond to the individual reference sequences shown in the plot legend.

### Viral *in situ* detection and transmission electron microscopy

Following metagenomic investigation, a coronaviral RNA probe was designed for RNAscope ISH targeting the nucleoprotein sequence aligning with the present coronavirus. Coronaviral RNA was abundantly detected in the small intestine (9/9, 100%, Figure 1E) and the caecal mucosa (6/6, 100%) of all individuals originating from the three selected cases (Suppl. material – S9). The positivity was identified in the cytoplasm of the enterocytes lining the villi of the small intestine (Figure 1E) and caeca, as well as subepithelial and sloughed luminal cells. Coronaviral RNA was also detected within caecal tonsils and bursal follicles. The rest of the tissues, including pancreas, trachea and kidney, of all birds examined were negative.

Finally, ultrastructural analysis was performed on well-preserved intestinal tissue sections from three birds exhibiting marked enteric lesions at histopathology and strong RNA detection by RNAscope ISH. Numerous round, membrane-bound, particles (90–100 nm in diameter) were observed within large vesicles in the cytoplasm of villous enterocytes of one bird (Figure 1F-H). These features were consistent with enveloped virions (28). No particles were detected in glandular enterocytes

## Discussion

This case series describes enteritis associated with a novel coronavirus in meat-type chickens, distinct from IBV. An important feature of the disease was the late onset of clinical signs compared to the more classical presentation of viral enteritis in poultry, which typically affects very young birds (7–42 days) (29). At this later stage of production, the economic consequences are greater, due to peak mortality as well as the risks of condemnation and downgrading at the processing plant.

The clinical signs and lesions observed closely resemble those described in so-called “fulminating disease” in guinea fowl, including chilliness and ruffled feathers (8). Macroscopically, enteritis and a “snowy” pancreas were similarly observed. Histopathology corroborated the necropsy findings, revealing severe villous atrophy and enterocytic cytopathy, strongly supporting the involvement of an epitheliotropic virus, which was subsequently confirmed by RNAscope ISH. In parallel, the contribution of coccidial infection was ruled out, as both mucosal scrapings and histology were negative in most cases and only positive in a single flock.

Overall, the predominance of coronaviral sequences in several independent cases, in the absence of other known parasitic, bacterial or viral agents, combined with the colocalization of coronavirus RNA within enteritis lesions, strongly supports the involvement of a novel coronavirus as the causal agent. A more robust validation of causality would require virus isolation assays and experimental infection studies.

To date, coronavirus-related enteritis in chickens has been associated exclusively with IBV, in both natural and experimental infections in 14- to 36-day-old broilers in the USA and Brazil (30-32). Several of the IBV strains involved were characterized and grouped as California enteric variants (CalEnt). In contrast to these previously described cases, which involved clinical signs or viral detection in the respiratory tract, our findings support an almost exclusive tropism for the digestive tract.

Interestingly, the coronavirus identified in our study clusters with GfCoV and, to a lesser extent, TCoV, both of which are known for their enteric tropism. As neither GfCoV nor TCoV has ever been recovered from chickens with enteritis, we tentatively named this viral lineage Chicken enteritis coronavirus (ChECoV). Although the overall genome organization (pp1a–pp1b–S–3a– 3b–E–M–4b–4c–5a–5b–N–6b) is consistent across the strains analyzed in this study, the absence of one accessory gene (6b) in one case appears to be strain-dependent. However, its presence is not required for the production of viable viral progeny, and its role in viral pathogenesis remains unclear (33,34). The presence of this auxiliary, apparently non-essential accessory gene reflects the genomic flexibility of AvCoV, which not only tolerates the stable insertion of novel genes but also allows genomic reorganization (35).

Fine-scale phylogenetic analyses and recombination breakpoint detection in avian enteric coronavirus genomes should be interpreted with caution, owing to the limited and biased representation of TCoV, GfCoV and other enteric AvCov sequences relative to IBV genomes. The latter are vastly overrepresented as a consequence of the intensive worldwide surveillance of infectious bronchitis in the poultry industry.

Our findings raise questions regarding the adaptation of the virus to the chicken host and, more specifically, the molecular determinants underlying this shift in tropism in GfCoV-related coronaviruses. So far, IBV, TCoV and GfCoV have shown distinct phenotypes in terms of host and tissue tropism, and experimental challenge with TCoV in chickens has failed to reproduce enteric lesions (36). Here, GfCoV spike genes show a striking evolutionary trajectory from 2011 to 2025, likely driven by a continuous immune pressure and that might explain the evolution of host tropism toward chickens and warrants further functional studies based on glycan or tissue binding assays: such a study previously suggested an evolution of GfCoV tissue tropism associated to a similar evolution on spike gene (6). The circumstances that led to this host-range expansion toward chickens remain unclear, but the practice of raising chickens and guinea fowl in close proximity (concurrently in the same farm or successively in the same barn), a common practice in this region of France, and very unlikely elsewhere, may have provided favorable conditions for viral adaptation between these avian species. This emphasizes the impact of farming practices (stocking densities, mixes of different species, contacts between domestic and wild animals, …) on the emergence of animal and zoonotic viruses (37).

Our findings also raise concerns on the reliability of “historical” vernacular naming of avian coronaviruses, mostly based on their host species, and the need for a standardized terminology (27). Additional research is needed to identify and characterize coronaviral receptors in host cells (6), as well as to investigate the origin of the virus, the occurrence of recombination events, and potential reservoirs.

The emergence of an enteropathogenic coronavirus in chickens may represent a highly significant challenge for the global broiler industry and warrants intensive research and surveillance.

## Acknowledgments

We are grateful to the Genotoul bioinformatics platform Toulouse Occitanie (Bioinfo Genotoul, https://doi.org/10.15454/1.5572369328961167E12) for providing help and/or computing and/or storage resources. We acknowledge the contribution of attending veterinarians for clinical investigations and sampling. The genomics analyses have been done in the framework of the METAPATH clinical metagenomics platform, with the financial funding of the Région Occitanie, France. We acknowledge the CMEAB, member of the national infrastructure France-BioImaging (https://ror.org/01y7vt929) supported by the French National Research Agency (ANR-24-INBS-0005 FBI BIOGEN). We sincerely thank the IT department of ENVT for their support with informatics infrastructure.

