## Supplementary material for "A novel coronavirus associated with enteritis in broiler chickens, France, 2025"

### S1. Methods

#### Recombination analysis

We looked for putative recombination events in the sampled genomes using the tool SHERPAS (1). Briefly, Sherpas builds a phylo-k-mer database from a phylogeny of typed viruses. Then, query genomes are scanned via a sliding window and compared to this database to extract a list of genome segments with associated types. To build the SHERPAS database, we used the tree generated in this study, complemented with genomes from (2) and (3) (pigeon and quail coronaviruses respectively). SHERPAS was run with default parameters to the exception of the phylo-k-mer length that was increased to k=11 (as recommended in original manuscript). Recombination pattern figures were generated with the R script present in SHERPAS repository.

#### Phylogeny and similarity analysis

Nucleotide sequences were aligned using MAFFT. Phylogenetic trees were reconstructed under the maximum likelihood framework using IQ-TREE, following the W-IQ-tree workflow (4). For the whole-genome dataset (WG), the best-fit substitution model selected by Model Finder GTR+F+R4 branch support was assessed using Ultrafast Bootstrap with 1000 replicated. For the S gene dataset, the best-fit model was TIM3+F+G4, with branch support similarly estimated using 1000 UFBoot replicated. To detect potential recombination events, similarity plots were generated using SimPlot++ with Kimura-2-parameter distance model, a window size of 200 nt and a size of 20 nt (5).

#### Transmission electron microscopy

Tissues stored in formalin were cut immersed in 2% glutaraldehyde in Sorensen's phosphate buffer (0.1 mol/L, pH = 7.4) for 1h, and washed with Sorensen's phosphate buffer for 12 h. Then samples were incubated with 1% $OsO_4$ in Sorensen's phosphate buffer (0.05 mol/L, glucose 0.25 mol/L, $OsO_4$ 1%) for 1 h, dehydrated in an ascending ethanol series until ethanol 100° and then with propylene oxide, and embedded with epoxy resin (EMBed 812). After 48h of polymerization at 60°C, ultrathin sections (70 nm) were mounted on 100 mesh collodion-coated copper grids and post-stained with 3% uranyl acetate in 50% ethanol and with 8.5% lead citrate before being examined on a HT 7700 Hitachi electron microscope at an accelerating voltage 80 KV.

**S2**. **Temporal dynamics of daily mortality in broiler flocks and respective barns.** Mortality data is given as a percentage and presented for each barn to capture barn-level dynamics. Cases 59/2025 and 239/2025 correspond to a single flock (same age, same origin) dispatched among several barns.

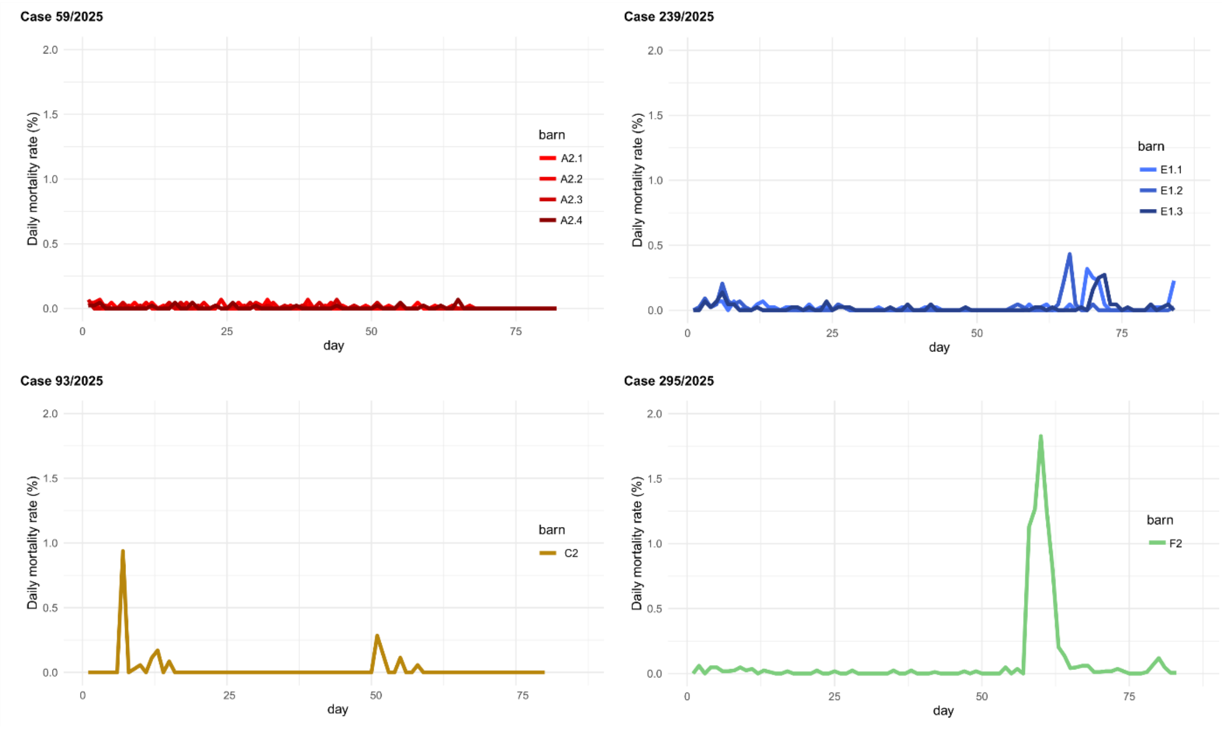

**S3. Clinical data from the different enteritis cases included in this study.** The estimated severity of the enteritis considers the estimated percentage of diseased birds in the flock along with the severity of the clinical signs and the lesions at necropsy. Mortality data includes only deaths occurring after the onset of enteritis. Slaughterhouse data on condemnations and downgrades were collected when available. No quantitative data could be obtained specifically for each type of condemnation reason.

| Lab ID | Clinical enteritis (onset of signs) | Enteritis-related mortality (%) | Condemnations (%) | Downgrades (%) |
|---|---|---|---|---|
| 59/2025 | Severe (60 days) | 0 | 1.3 | 7.5 |
| 93/2025 | Mild (51 days) | 1.4 | 1 | 4.6 |
| 239/2025 | Severe (70 days) | 0.8 | 0.6 | 7.4 |
| 295/2025 | Severe (60 days) | 6.9 | 2.5 | - |

**S4. Prevalence of histopathological findings detected in tissue sections from broiler chickens naturally infected with ChECoV (4 flocks)**. For each tissue and flock, histopathological findings are expressed as the proportion of affected birds to the total number of birds examined. NA: not available.

| Flocks | 239/2025 | 59/2025 | 295/2025 | 93/2025 | Total (%) | Histopathological findings |
|---|---|---|---|---|---|---|
| Small intestine | 7/8 | 4/4 | 6/6 | 2/4 (1) | 19/22 (86%) | Villous atrophy, fusion and blunting with enterocyte and subepithelial necrosis, infiltration of lamina propria by lymphocyte-rich leukocytes. |
| Pancreas | 7/8 | 4/4 | 6/6 | 2/4 | 19/22 (86%) | Pancreatic acinar cell atrophy with zymogen granules depletion, vacuolization, single cell necrosis/apoptosis. |
| Thymus | 1/8 | 3/4 | 4/5 | 1/2 | 9/14 (64%) | Diffuse thymic atrophy with cortical lymphocytic depletion. |
| Bursa | 3/8 | 1/4 | 5/5 (2) | 1/4 | 10/21 (48%) | Diffuse bursal atrophy with lymphocytic depletion, epithelial and/or follicular cysts, hyalinosis and epithelial folding |
| Gizzard | 1/8 | NA | NA | 1/4 | 2/12 (17%) | Erosion and ulceration of gizzard mucosa with fibrino-heterophilic exudate and intralesional bacteria |
| Trachea | 2/8 | NA | NA | 0/4 | 2/12 (17%) | Diffuse lymphocytic tracheitis (2/12), with fibrino-heterophilic tracheitis and intraluminal bacteria (1/12). |
| Proventriculus | 0/8 | 2/4 | NA | 0/4 | 2/16 (13%) | Focal to extensive epithelial necrosis and sloughing of mucosal proventricular glands. |
| Lung | 1/8 | NA | NA | 0/4 | 1/12 (0,8%) | Exudative fibrino-heterophilic primary bronchitis with intraluminal bacteria (cocci). |
| Kidney | 0/8 | 0/4 | 0/6 | 0/4 | 0/22 (0%) | - |
| Spleen | 0/8 | 0/4 | 0/6 | 0/4 | 0/22 (0%) | - |

(1) Intestinal coccidiosis (4/4); (2) Bursal cryptosporidiosis (4/5).

**S5. Sequencing metrics for the four samples processed using the metagenomics workflow**.

| Case | 59/2025 | 93/2025 | 239/2025 | 295/2025 |
|---|---|---|---|---|
| **Host** | Chicken | Chicken | Chicken | Chicken |
| **Sample type** | Intestine | Intestine | Intestine | Cloacal swab |
| **Ct value (Maurel 2011)** | 12.78 | 16.46 | 7.39 | 9.3 |
| **Total bases (Mb)** | 1,793.82 | 2,900.81 | 8,280.24 | 130.46 |
| **Passed bases (%)*** | 89.3 | 91.6 | 89.68 | 82.1 |
| **Total reads** | 4,527,320 | 3,695,240 | 15,809,440 | 233,230 |
| **Passed reads (%)*** | 89.1 | 91.8 | 90.6 | 81 |
| **Filtered reads **** | 357,062 | 1,406,528 | 237,073 | 19,964 |
| **Viral reads** | 90,208 | 113,518 | 49,909 | 14,819 |
| ***Gammacoronavirus* reads** | 72,281 | 5,196 | 40,595 | 13,324 |
| **Accession number** | PV843471.1 | PV843472.1 | PX987256.1 | PV843474.1 |

*Reads/bases must have a quality score above 9 to pass.

**Reads must be larger than 500bp, not map on the *Gallus gallus* genome and have a quality score above 15 to pass.

**S6. Representative taxonomic assignment of viral and bacterial reads from metagenomic analysis conducted on the intestinal samples from affected chickens (case 59/2025).** Kraken2 classification using the bacterial/viral/archaea database identified 267,727 bacterial reads.

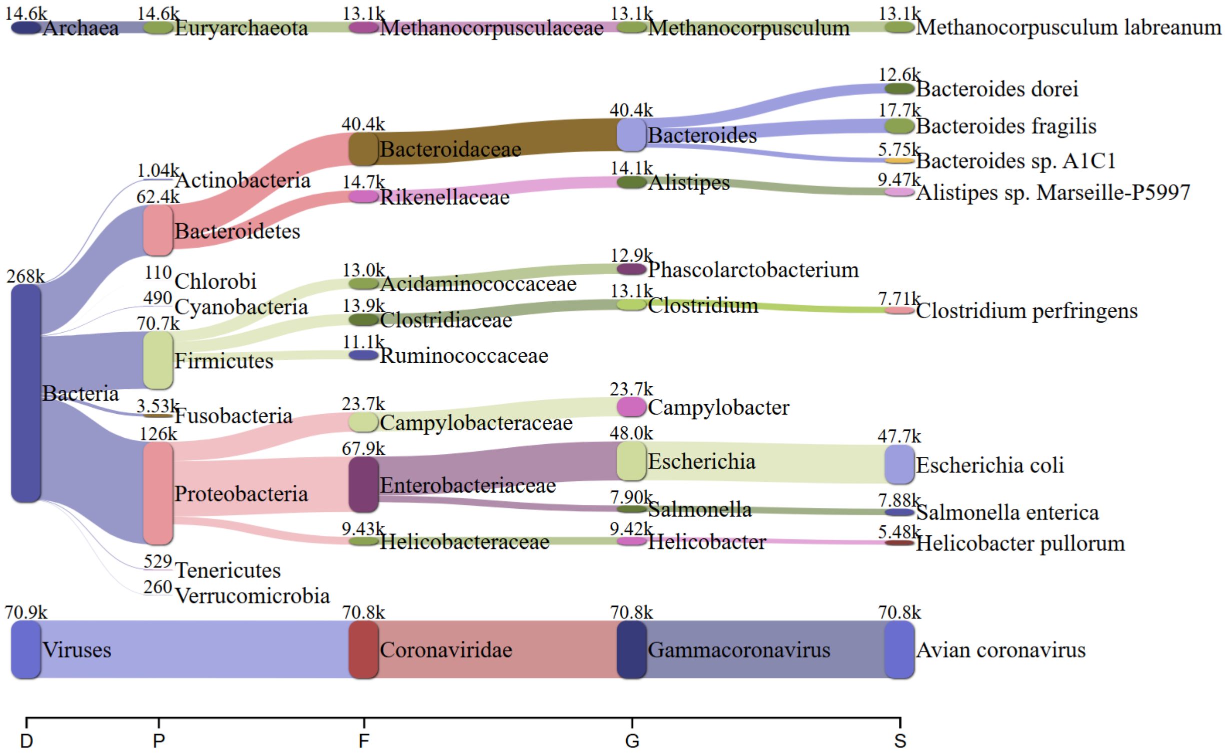

**S7.** Results (Ct values) of real-time gammacoronavirus RT-PCR (Maurel et al, 2011) performed on intestinal content, cloacal swabs and environmental wipes, on the four included cases. All tested samples were positive, empty table cells do not correspond to a negative result.

| Lab ID | *Gammacoronavirus* RT-PCR detection [Ct value range] | | |
|---|---|---|---|
| | **Intestinal content** | **Cloacal swabs** | **Dust (wipes)** |
| 59/2025 | [12.78] | - | - |
| 93/2025 | - | [16.46 - 24.42] | [19.23 - 21.52] |
| 239/2025 | [7.39 - 15.07] | - | - |
| 295/2025 | [9.30- 24.17] | [16.19 - 18.60] | [26.26] |

**S8. Recombination patterns predicted in the four sample genomes.** Each genome (Y-axis) is decomposed into intervals which are defined by their genome coordinates (X-axis, expressed in base pairs) and which are assigned to their most likely taxa of origin from the reference tree. Taxa and corresponding color code are described in the legend. All genomes show a phylogenetic signal matching the GfCoV taxon in the Spike locus, which is located between positions 20,000 and 25,000. Following the SHERPAS tools author's recommendation, only the large intervals accounting for more than 300bp (the parametrized window size) should be considered as a putative recombination signal. Shorter intervals can be due to spurious local homologies favored by taxon over-representation in the reference tree, which is indeed the case here for IBV_Mass-like and IBV_QX-like taxa. N/A intervals represent regions that could not be assigned with confidence to any taxa.

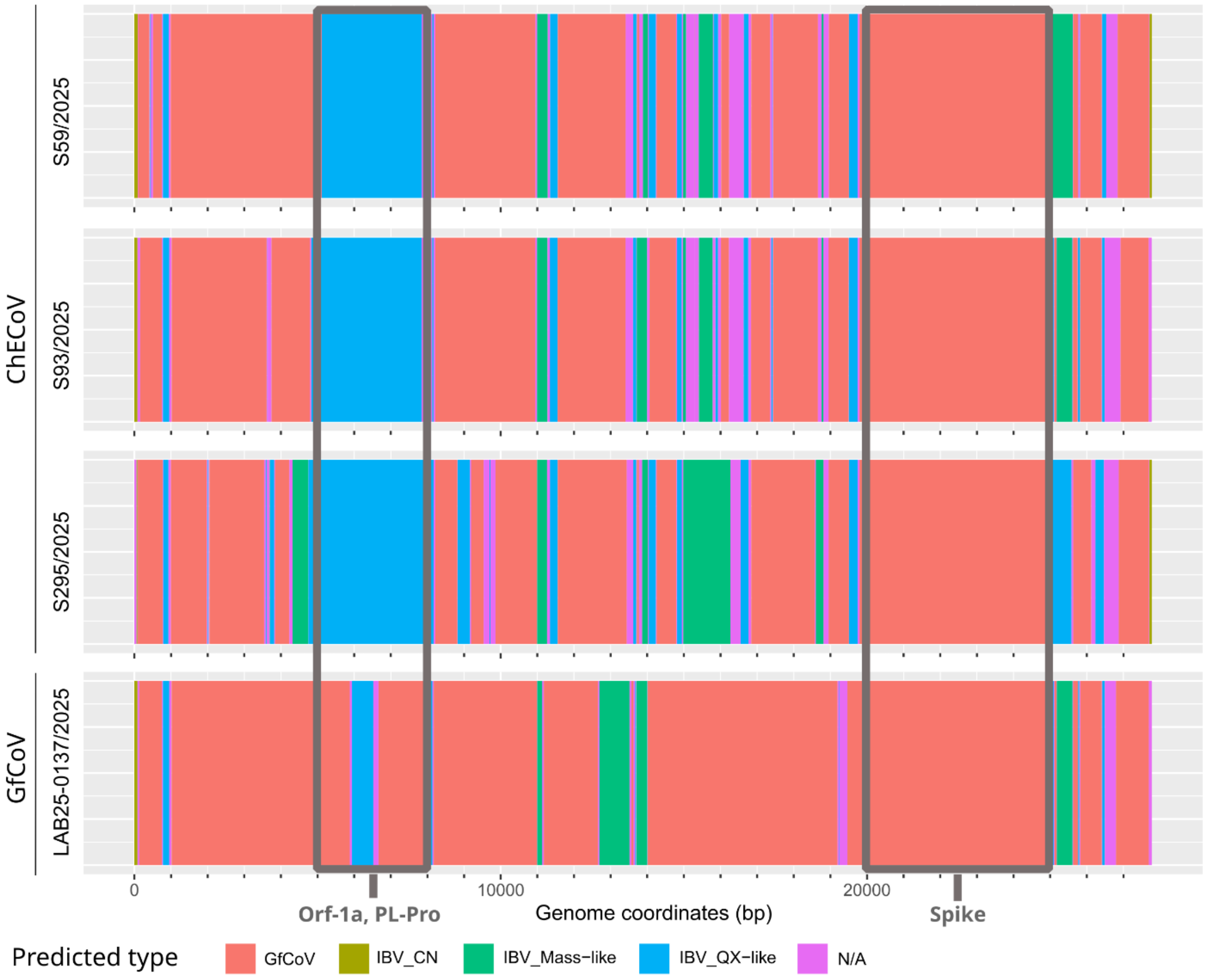

**S9.** Detection of gammacoronavirus RNA in tissue sections from chickens naturally infected with ChECoV by RNAscope *in situ* hybridization (3 flocks, and 3 birds/flock). For each tissue and flock, histopathological findings are expressed as the proportion of affected birds relative to the total number examined. The mean semi-quantitative score is indicated in brackets (0= none; 1=sparse; 2=frequent; 3=widespread. NA: not available.

| **Flocks** | **239/2025** | **59/2025** | **295/2025** | **Total** | **Main positive cell types and structures** |
|---|---|---|---|---|---|
| Small Intestine | 3/3 (3) | 3/3 (3) | 3/3 (2) | 9/9 (3) | villous enterocytes, sloughed cellular debris within lumen |
| Caeca | 3/3 (3) | NA | 3/3 (3) | 6/6 (100%3) | apical enterocytes, mucosal lymphoid follicles, sloughed cellular debris within lumen |
| Bursa | 1/3 (1) | 2/3 (1) | 0/2 (0) | 3/8 (1) | sparse cells within few bursal follicles |
| Pancreas | 0/3 (0) | 0/3 (0) | 0/3 (0) | 0/9 (0) | - |
| Thymus | 0/3 (0) | 0/3 (0) | 0/3 (0) | 0/9 (0) | - |
| Trachea | 0/3 (0) | NA | NA | 0/3 (0) | - |
| Proventriculus | NA | 0/3 (0) | NA | 0/3 (0) | - |
| Lung | 0/3 (0) | NA | NA | 0/3 (0) | - |
| Kidney | 0/2 (0) | 0/3 (0) | 0/3 (0) | 0/8 (0) | - |
| Spleen | 0/3 (0) | 0/3 (0) | 0/3 (0) | 0/9 (0) | - |
